# Enhanced Proteomics Analysis with a Novel Recombinant Chymotrypsin Analogue Engineered for High Cleavage Specificity

**DOI:** 10.64898/2025.12.19.695449

**Authors:** Kish R. Adoni, Jonathan E. Ditcham, Alba Katiria González Rivera, Georgina H. Charlton, Sergei Saveliev, Konstantinos Thalassinos, Riccardo Zenezini Chiozzi

## Abstract

Chymotrypsin is widely used in shotgun proteomics owing to its orthogonal cleavage specificity relative to trypsin, which enhances sequence coverage of hydrophobic protein regions. However, commercial preparations often display variable cleavage specificity, trypsin contamination, and elevated missed-cleavage rates, which can collectively reduce proteome coverage and data reproducibility. To address these limitations, we present a novel recombinant chymotrypsin (*rChymoSelect*) engineered for improved cleavage specificity and robustness in proteomics workflows. Benchmarking against standard bovine chymotrypsin revealed 97 % C-terminal cleavages after tyrosine (Y), phenylalanine (F), and leucine (L) for *rChymoSelect*, compared with 72 % for the standard enzyme. This enhanced cleavage specificity reduced missed cleavages and increased peptide-spectrum matches across charge states. Across 3,638 identified proteins, *rChymoSelect* yielded 22.2 % unique identifications compared with 8.2 % for standard chymotrypsin, while maintaining similar peptide length, m/z, and hydrophobicity distributions. Notably, *rChymoSelect* showed enriched recovery of mitochondrial proteins, consistent with its improved digestion of hydrophobic targets. The enzyme remained active in up to 6 M urea and achieved near-maximal proteome coverage within 2 hours (only a 2.4 % gain after overnight digestion). Integration with data-independent acquisition (DIA) increased total protein identifications from approximately 2,200 (DDA) to 3,200 (DIA), a 45 % gain, with *rChymoSelect* outperforming standard chymotrypsin by 16.6–22.4 % in peptide-spectrum matches and 4.6–6.2 % in protein identifications. These results establish *rChymoSelect* as an advanced tool with improved cleavage specificity that reduces analytical complexity and enhances the reliability of proteomic analysis, while expanding chymotryptic digestion to hydrophobic and high-denaturant proteomics applications.

## Introduction

Recent advancements in sample preparation methodologies, liquid chromatography, mass spectrometry and data processing workflows have spearheaded a revolution in the field of proteomics, leading this methodology to represent the gold standard for high-throughput protein identification^1,2^. During proteomics sample preparation, trypsin is commonly used as the enzyme of choice for the conversion of proteins into appropriately charged and sized peptides for subsequent proteomics mass spectrometry analysis. Since trypsin cleaves specifically at the C-terminus of lysine and arginine, there are some limitations to its utility for protein analysis, particularly in the context of hydrophobic proteins, membrane proteins, proteins with excessive lysine/arginine amino acids in the primary sequence and proteins with excessive post-translational modifications at lysine/arginine residues^3^.

To this end, chymotrypsin provides a powerful orthogonal approach towards proteomics sample preparation^4^. Whilst both proteases share the same serine-protease catalytic triad, via serine (S195), histidine (H57) and aspartic acid (D102), their differences are driven by the differential substrate binding pocket (S1), with the negatively charged aspartic acid of trypsin’s S1 pocket facilitating hydrolysis at the C-terminus of lysine (K) and arginine (R). Conversely, the deeply hydrophobic nature of chymotrypsin’s S1 binding pocket facilitates its preference for cleavage at larger, hydrophobic residues such as phenylalanine (F), tyrosine (Y) and tryptophan (W)^5^. Consequently, chymotrypsin targets the C-terminus of residues: phenylalanine, tryptophan, tyrosine, methionine (M) and leucine (L). As such, these properties have been leveraged for more comprehensive hydrophobic protein digestion^6,7^. Common limitations of chymotrypsin in the context of proteomics include its promiscuity of cleavage specificity and the increased occurrence of missed cleavages^8^. Furthermore, chymotrypsin is traditionally derived from bovine pancreatic tissue, potentially leading to contamination with trypsin in commercial products. Several manufacturers note this explicitly, as some commercial formulations are treated with TLCK to inhibit trypsin activity^9^. The increased heterogeneity of this digested peptide population, from chymotrypsin digestion, can greatly hinder the data processing step of proteomics experiments, on account of increased computational search space requirements, leading to reduced and even false peptide identifications^10^.

To address these limitations, we evaluated a novel recombinant chymotrypsin (rChymoSelect, Promega) engineered for improved cleavage activity and specificity at Y, F and L, to reduce off-target amino acid site cleavages and missed cleavages. Further, rChymoSelect is calcium independent, resistant to autoproteolysis and maintains activity under mildly denaturing conditions (*Promega-reference or data to demonstrate this*). In this work, we benchmarked rChymoSelect against standard chymotrypsin, in the context of high-throughput proteomics. Our findings demonstrate that rChymoSelect’s enhanced cleavage specificity boosts efficiency and reproducibility for modern proteomics workflows, leading to increased protein identifications and protein sequence coverage, relative to standard chymotrypsin, without affecting the physiochemical properties of the digested peptides.

## Materials and Methods

### rChymoSelect production

rChymoSelect is a recombinant chymotrypsin under development by Promega. The enzyme was provided at a concentration of 1 µg/µL and stored at –80°C. Prior to use, aliquots were thawed on ice and used without further preparation.

### Protein preparation

For all experiments relating to the benchmarking of rChymoSelect against standard chymotrypsin (V1061, Promega), a standard human K562 intact extract (10mg/mL of human cell K562 lysed in 6 M Urea, Promega V6941) was used. For comparison of rChymoSelect to trypsin (Trypsin Gold, V5280, Promega); human bone osteosarcoma epithelial (U2-OS) cells were lysed in 100 mM Tris-Cl (pH8.5), 5% sodium dodecyl sulfate (SDS), 5mM Tris(2-carboxyethyl)phosphine hydrochloride (TCEP) and 20mM chloroacetamide. Samples were boiled for 10 minutes and then cooled to room temperature before being subjected to ultrasonic bath sonication for 10 minutes. Protein content was quantified via Bicinchoninic Acid (BCA) assay^11^ (ThermoFisher Scientific).

### Optimisation of in-solution and SP3 digestion

20µg of K562 lysate, was diluted to a final concentration of 2M, 4M and 6M urea for rChymoSelect, 2M and 0.6M urea for standard chymotrypsin and 0.6M urea for trypsin. All digestions were performed at 25°C for in-solution digestion, at 2- or 16-hours using a thermoshaker (Eppendorf) at 1500rpm. For SP3 digestion, the samples were prepared in a KingFisher APEX robot (ThermoFisher Scientific) using a protocol from Koenig et al^12^, with the following modifications: the 96-well comb was stored in plate 1, plate 2 contained reconstituted sample in 70% acetonitrile with magnetic MagReSyn Hydroxyl beads in a protein/bead ratio of 1:2. Washing solutions were in plates 3–5 (95% acetonitrile) and plates 6–7 (70% ethanol). Plate 8 contained 200 μL of digestion solution, which consisted of protease (rChymoSelect or standard chymotrypsin) in 50 mM ammonium bicarbonate (pH 8.5). For this optimisation, different protease/protein ratios (1:100, 1:40, 1:20) and guanindinium hy-drochloride (GND) concentrations (0.1M, 0.2M and 0.4M) were tested, with incubation for 2-hours, at 37°C. For both digestion protocols, protease activity was quenched by acidification with trifluoroacetic acid (TFA) to pH 2 before the resulting peptide-mixture was desalted on an OASIS HLB 96-well plate (Waters), before drying *in-vacuo* using a Savant DNA120 (ThermoFisher Scientific).

### Comparison of Trypsin and rChymoSelect with deep offline fractionation

200 µg of proteins from U2-OS cells lysed in SDS buffer were prepared using the SP3 lysis method and digested with either trypsin overnight or rChymoSelect for 2h at 37 °C. Digestion was quenched by TFA, and peptides were desalted using the OASIS HLB 96-well plate (Waters). Peptides were then fractionated with a Vanquish HPLC (Thermo Fisher Scientific) using an Acquity BEH C18 column (2.1 x 50 mm with 1.7µm particles, Waters), mobile phase A: 10 mM ammonium formate, pH 10, mobile phase B: 80% acetonitrile, 10mM ammonium formate pH10, with a flow rate of 1000µL/min. 24 fractions were collected across a 6-minute gradient. Fractions were then dried and reconstituted in 0.5% TFA before undergoing liquid chromatography–tandem mass spectrometry (LC-MS/MS) analysis.

### LC-MS/MS

Peptides were reconstituted in 0.5% TFA, and 500ng were injected on an UltiMate 3000 RSLCnano liquid chromatography system (Thermo Fisher Scientific). A 5.5 cm μPAC Neo HPLC analytical column (COL-CAPHTNEOB) was connected to a Silica Tip emitter. Column temperature was set at 45°C, and column flow rate was set to 1500 nL/min. Mobile phase A (0.1% formic acid) and mobile phase B (80% acetonitrile, 0.1% formic acid) were applied with an elution gradient from 1.0 to 35.0% mobile phase B over either 25 minutes (for the fractionations) or 51 minutes (protocol optimization). Peptides were ionized using a spray voltage of 2.0 kV (275°C). The mass spectrometer was set to acquire full-scan MS spectra (350 to 1400 m/z) for a maximum injection time set to Auto at a resolution of 120,000 and an automated gain control (AGC) target value of 250%. The instrument was set up in DDA with a Wide Window Acquisition (4th window) for all the precursors with a charge state in range of 2+ to 5+; most abundant ions were then fragmented in HCD (30% normalized collision energy) with MS2 acquisition at a resolution of 30,000, AGC: 400%, maximum injection time: Dynamic, dynamic exclusion: 20s. For protocol optimization with Data-Independent Acquisition: 56 windows of 12 Da (*m/z* range 361-1033) were fragmented via HCD with normalized energy: 30%, AGC: 1000% and resolution: 15,000.

### Data Analysis

MS data were searched against the Human SWISS-Prot protein database (August 2023) with SAGE proteomics software for downstream analysis^13^ (for DDA). The increased capacity to handle large search space (due to increased cleavage sites and missed cleavages), as well as its fast performance made it an appropriate software choice for this work^14^. For cleavage specificity analysis, the following parameters were set: cleavage sites: F, Y, W, L, K, R, H, N, Q (except before P), missed cleavages: 6. For onward analysis, cleavage sites: FYWL (except before P), missed cleavages 3. For all searches, variable modifications: [+15.9949, M]; static modifications: [+57.0215, C], max variable modifications: 3. PSMs with q-value<0.01 were selected for onward analysis, ambiguous proteins and contaminants were filtered out. Further, all validated proteins were identified in at least 2 of 3 biological replicates. Raw SAGE search outputs were processed with the PickedGroupFDR pipeline^15^ to produce a combined_protein.tsv file that was used for protein quantification and downstream enrichment analyses. Peptide-protein rollup and proteingroup collapsing were performed using the PickedGroupFDR implementation. Downstream statistics, plotting and figure generation were performed in GraphPad Prism^16^. For Data-Independent Analysis (DIA), RAW data was processed with FragPipe v23.1^17^, with enzyme: chymotrypsin cleavage sites: F, L, W, Y, (except before P), missed cleavages: 1, variable modifications: [+15.9949, M], [+42.0106, Nt], fixed modifications: [C, +57.02146] and modifications per peptide: 3. For DIA library-based searches, the DDA RAW files from the corresponding runs were used to generate a spectral library, using DIA-NN^18^ Unless otherwise stated, all samples were analysed in experimental triplicate, and all mass spectrometry data is available via Proteome Xchange (PRIDE accession: PXD072165).

## Results and Discussion

We performed a comprehensive characterisation of rChymoSelect against its commercially available analogue (defined here as standard chymotrypsin), as well as investigating the optimal digestion conditions in the context of buffer constitution, digestion methodology and incubation time (Fig. 1).

**Fig. 1:**
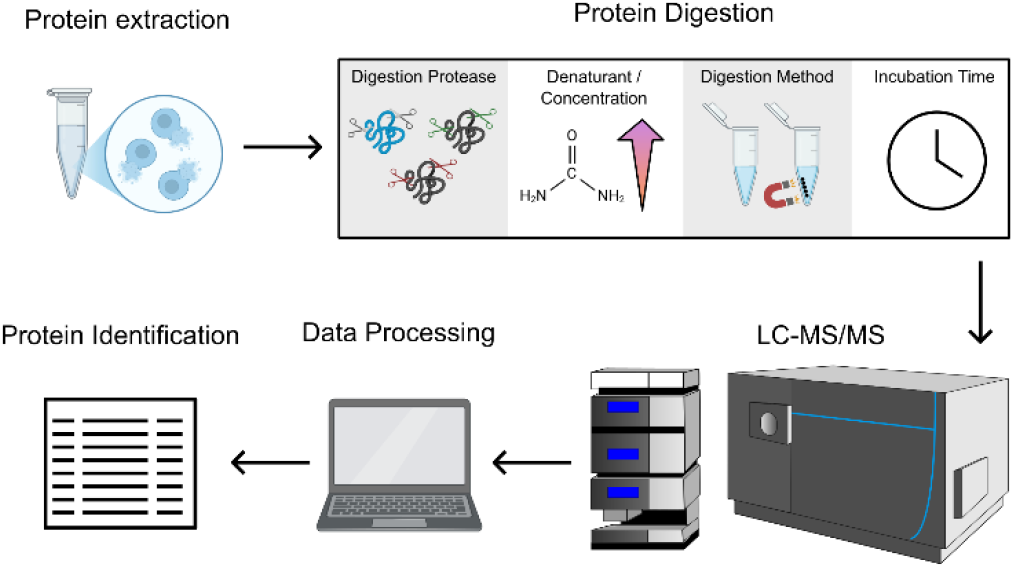
Benchmarking workflow for rChymoSelect vs standard chymotrypsin. Digestion enzyme (rChymoSelect vs chymotrypsin), buffer denaturant concentration (urea: 2M, 4M, 6M, guanidinium hydrochloride: 0.1M, 0.2M, 0.4M), digestion methodology (in-solution vs SP3) and incubation time (2h vs 16h) were all investigated against intact human protein lysate (Promega, V6941).

### rChymoSelect boosts cleavage specificity and reduces missed cleavage identifications

Standard chymotrypsin’s promiscuity in the context of cleavage site specificity can impede downstream proteomics data processing pipelines. This occurs due to the increased search space that is associated with the inclusion of up-to-9 amino acid cleavage sites, as well as the increased number of missed cleavages that must be accounted for in the processing software. Consequently, the larger computational burden can drive slower data processing times, and perhaps more importantly, an increase in false positive identifications^10^. Two computational approaches can be used to curtail this cleavage site heterogeneity. Firstly, one can include only the most probable cleavage sites (L, F, Y) and ignore all peptides that were generated from other “off target” cleavages. However, this approach neglects these non-L, -F, and-Y peptides from the theoretical candidate peptide library for onward peptide spectral match (PSM) identification. Alternatively, all 9 possible cleavage sites can be incorporated into the data processing workflow, to ensure that all peptides can theoretically be identified. Whilst this facilitates the identification of more PSMs, the pitfalls that are associated with this exponential increase in computation search space requirements are incorporated into the results.

To investigate whether cleavage specificity was improved with rChymoSelect, we initially performed a chymotrypsin proteomics experiment against commercially available intact protein lysate from human K562 cells (Promega) (Fig. 1). Comparison of both digestion enzymes at 2-hour and 16-hour incubation, with 2 M Urea, revealed a significant improvement in cleavage specificity for rChymoSelect vs standard chymotrypsin. Standard chymotrypsin analysis revealed that Leucine, Phenylalanine and Tyrosine cleavage sites accounted for just 72% and 82%, for 2 hour and 16-hour incubations, respectively. The remaining cleavage sites of lysine, arginine, tryptophan, histidine, asparagine and glutamine accounted for 28% and 17% of cleavages, for 2-hour and 16-hour incubation, respectively. In contrast, in the case of rChymoSelect, LFY site cleavage accounted for 97% and 95% for 2-hour and 16-hour incubation, respectively (Fig. 2a). The greatly improved cleavage specificity of rChymoSelect, whereby nearly all identified peptides were cleaved at L, F or Y, enables the user to incorporate just three cleavage sites with minimal loss of peptides, reduced computational burden and false positive identifications.

**Fig. 2:**
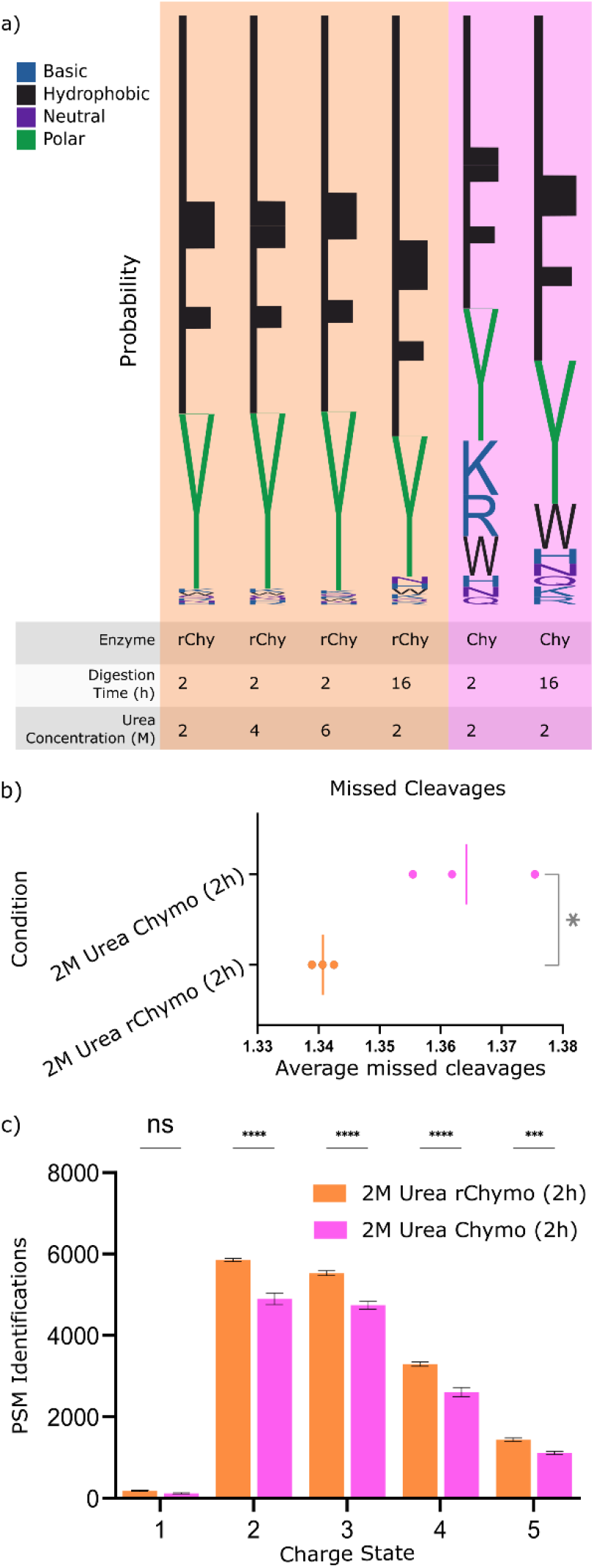
rChymoSelect enhanced cleavage specificity to reduce missed cleavage occurrence and increase protein identifications, vs standard chymotrypsin. (a) Cleavage site specificity analysis for rChymoSelect (rChymo) (digestion time: 2 h, 16h; urea concentration: 2M, 4M, 6M) vs standard chymotrypsin (Chymo) (digestion time: 2h, 16h; urea concentration: 2M). Cleavage sequence analysis performed using Logomaker^20^. (b) Number of missed cleavages for three experimental replicates of Chymotrypsin vs rChymoSelect (2M urea, 2h digestion). Statistical analysis performed using 2-sided unpaired t-test, * represents p<0.05. (c) Bar chart to represent PSMs for different charge states, rChymoSelect vs standard Chymotrypsin (2M urea, 2 h incubation).

As well as characterizing the improved cleavage specificity of rChymoSelect, we probed the optimal urea concentration for sample preparation. Urea represents a useful buffer component for proteomics sample preparation, due to its protein denaturing capacity. As such, more protein residues are exposed to the enzyme for digestion. Our findings suggested minimal changes in cleavage specificity of rChymoSelect at 2M-, 4M- and 6M-urea. Importantly, the application of chymotrypsin (and most proteases), is not amenable to high concentrations of chaotropic agents^19^. For example, urea concentration must be reduced to <1M in most currently available digestion protocols. These data demonstrate the effective functionality of rChymoSelect well beyond this threshold, suggesting its utility for digesting hydrophobic proteins that require aggressive denaturation conditions.

We next probed how the improved specificity of rChymoSelect could reduce the generation of missed cleavage peptides (Fig. 2b). Not only did rChymoSelect reduce the average number of missed cleavages, vs standard chymotrypsin, it also boosted the reproducibility across three experimental replicates. Reduced missed cleavage occurrence can boost peptide and protein identifications, as more peptides fit within the pre-specified precursor mass range of the quadrupole mass filter for precursor ion identification. Further, the maximum number of missed cleavages parameter can be reduced within the data processing workflow, again reducing computational burden and subsequent false-positive identifications. The increased reproducibility of cleavages also boosts the number of peptide-spectral matches (PSMs) for a given peptide, with positive consequences on peptide confidence and quantitation statistics. Our analysis revealed rChymoSelect generated more PSMs corresponding to 2+, 3+, 4+ and 5+ charges, relative to standard chymotrypsin.

Ultimately, rChymoSelect demonstrated reduced heterogeneity of cleavage sites, reduced missed cleavages and increased reproducibility of PSMs. These features combinatorially lead to greater peptide, and consequently, protein identifications.

### rChymoSelect does not alter the chymotryptic digested peptide physiochemical properties, relative to standard chymotrypsin

Having demonstrated the improved cleavage specificity and reproducibility of rChymoSelect, vs standard chymotrypsin, we next investigated whether these changes would modify the properties of the resulting chymotryptic peptides. No significant changes in peptide length were identified from our comparison across 2M urea, 2h incubations (Fig. 3a, b). Comparison of PSM *m/z* distribution revealed no significant increase, from rChymoSelect to standard chymotrypsin (Fig. 3c, d). PSM hydrophobicity, as measured by Grand average of hydrophobicity for a peptide or protein (GRAVY), also revealed no significant changes in peptide hydrophobicity from rChymoSelect, compared to standard chymotrypsin (Fig. 3e, f). These findings suggested that rChymoSelect enables improved digestion efficiency and reproducibility, leading to increased peptide and protein identifications, without modifying the physiochemistry of the peptides that were generated during sample preparation.

**Fig. 3:**
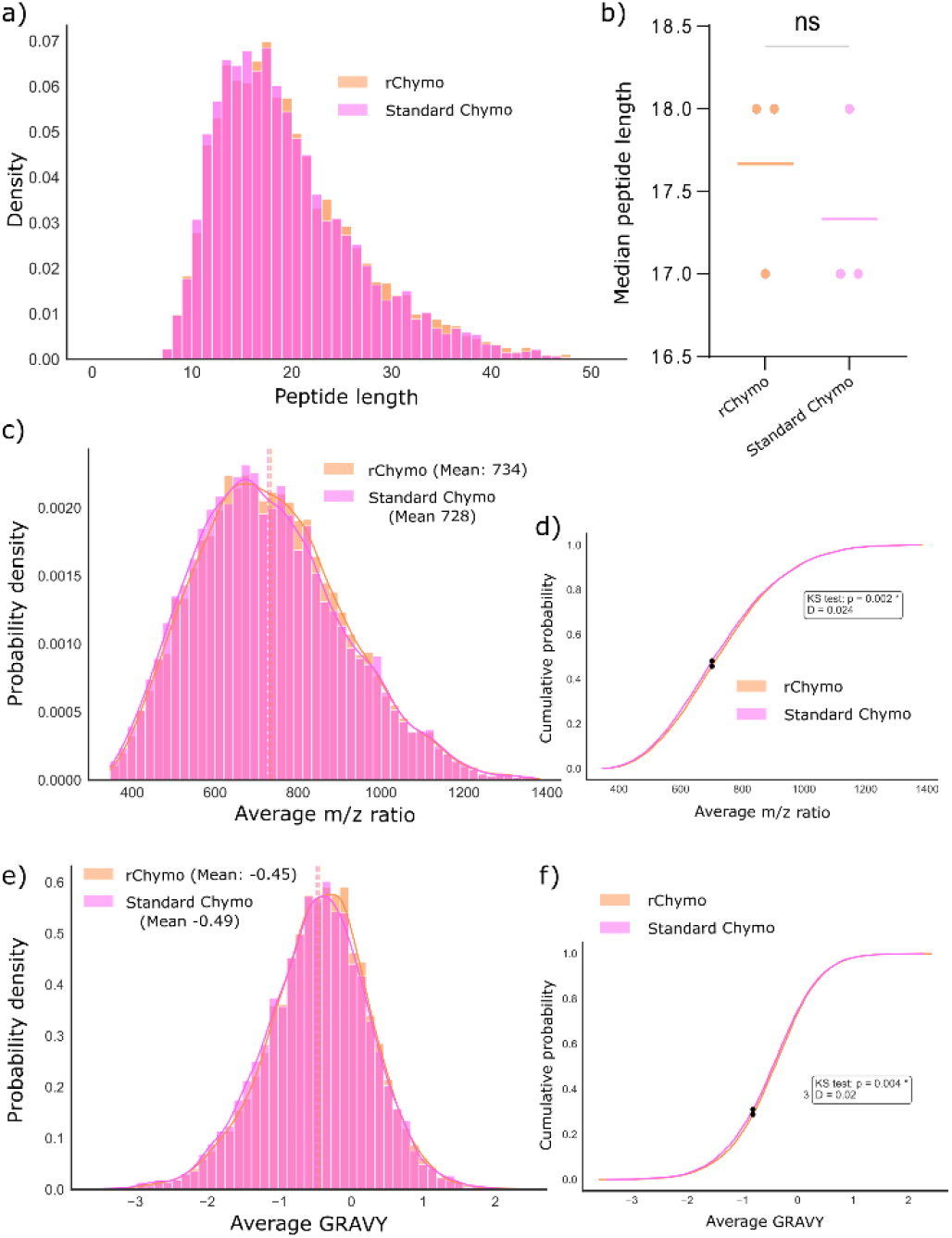
rChymoSelect does not alter peptide properties, vs standard chymotrypsin. (a, b) Comparison of peptide length between two proteases, as plotted by (a) density plot and (b) scatter plot. Statistical analysis performed using 2-sided unpaired t-test, ns represents p>0.05. Comparison of PSM *m/z* ratios as plotted by (c) probability density and (d) cumulative probability. Statistical analysis performed using two-sided Kolmogorov-Smirnov test for goodness of fit. (e, f) Grand average of hydrophobicity for a peptide or protein (GRAVY) score distribution as a measure of PSM hydrophobicity for each condition, demonstrated by (e) probability density and (f) cumulative probability. Statistical analysis performed using KS test, D=maximum absolute difference.

### rChymoSelect significantly boosts protein identifications, relative to standard chymotrypsin

Our benchmarking had thus-far revealed that rChymoSelect demonstrated improved cleavage specificity and reduced missed cleavage occurrence, without concomitant changes in peptide physiochemistry. These improvements led to a boost in PSM identifications across the standard proteomics precursor charge state distribution, relative to standard chymotrypsin. We next sought to investigate how these technical advancements in chymotrypsin activity would deliver more informative proteomics data. For this, we investigated the overall protein sequence coverage distribution for rChymoSelect vs standard chymotrypsin, across the proteome of human K562 cells (Fig. 4a). rChymoSelect demonstrated greater protein sequence coverage across the proteome, from the most abundant to least abundant identified protein of the dataset.

**Fig. 4:**
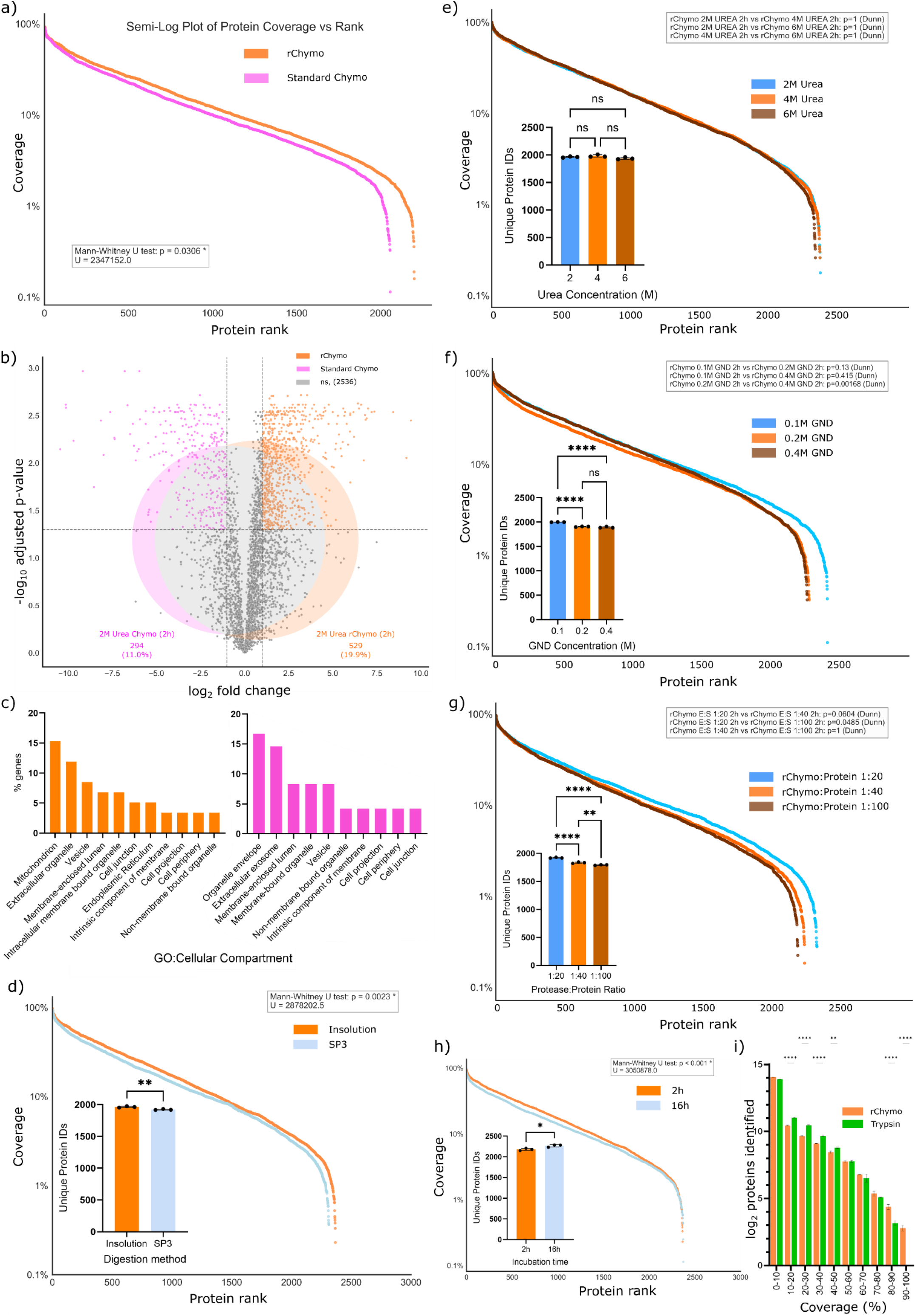
rChymoSelect significantly boosts protein identifications, relative to standard chymotrypsin, based on (a) protein sequence coverage distribution and (b) volcano plot and protein identifications Venn diagram. (c) Analysis of protein cellular compartment bias for rChymoSelect vs standard chymotrypsin, as depicted by bar chart of the most enriched gene ontology:cell compartment (GO:CC) (% genes/all annotated genes of GO:CC). All experiments were performed with 2M urea and 2-hour incubation using insolution digestion. Comparison of different sample preparation methodologies, including sequence coverage distribution and protein identifications bar-chart for (d) in-solution vs SP3 digestion (2M urea, 2 hour digestion, 1:40 rChymoSelect/protein ratio), (e) urea concentration (in-solution digestion, 2-hour incubation, 1:40 rChymoSelect/protein ratio), (f) guanidinium hydrochloride (GND) concentration (SP3 digestion, 2-hour incubation, 1:20 rChymoSelect/protein ratio), (g) rChymoSelect to protein ratio (SP3 digestion, no denaturant, 2-hour incubation), and (h) 2-hour vs 16-hour incubation (in-solution digestion, 2M urea, 1:40 rChymoSelect/protein ratio). (i) Protein identifications at different sequence coverages, across 10% sequence coverage bins, with comparison between rChymoSelect and trypsin using lysed U2OS cells, with offline peptide fractionation. Protein sequence coverage distribution plot statistical analysis performed using 2-sided Mann-Whitney U rank test on two independent samples, U represents the Mann-Whitney statistic. Volcano plot statistical analysis performed using 2-sided independent t-test, with Benjamin-Hochberg adjusted p-values. Bar chart statistical analysis performed using two-sided Kruskal-Wallis test with Dunn’s post-hoc test. rChymo vs trypsin bar chart statistical analysis performed using 2-way ANOVA, **represents p-value<0.01 and *** represents p-value<0.0001.

We next probed the differential proteomics data for rChymoSelect vs standard chymotrypsin. Of the 3638 proteins that were identified in our benchmarking experiment, 22.2% were exclusively identified from rChymoSelect, compared to 8.2% for standard chymotrypsin. These findings demonstrated the improved performance of rChymoSelect in generating a comprehensive proteomics dataset, vs standard chymotrypsin (Fig. 4b). Investigation of the cellular compartment of proteins that were identified from each proteolytic enzyme revealed broadly similar profiles. A notable exception was the significant enrichment of mitochondrial proteins from the rChymoSelect digestion, relative to standard chymotrypsin. This discrepancy between the two proteases could be explained by the relative increase in protein hydrophobicity for specifically mitochondrion localised proteins relative to other cellular compartments^21,22^. Furthermore, previous work has demonstrated that mitochondrial protein encoding genes tend to have greater hydrophobicity than nuclear protein-coding genes, potentially driven by the propensity for such hydrophobic domains to be recognized for endoplasmic reticulum associated degradation (ERAD), should they have been synthesized in the cytosol following nuclear gene transcription^23^. Consequently, the improved performance of rChymoSelect may have superior suitability for the study of mitochondrial proteins.

Following our benchmarking experiments for rChymoSelect vs standard chymotrypsin, we next investigated the optimal conditions, across our proteomics sample preparation workflow, to enhance rChymoSelect performance. Minimal differences in protein identifications were identified between in-solution vs SP3 digestion (2.15% more identifications for in-solution, Fig. 4d). However, automated SP3 digestion, using the KingFisher robot, provides multiple advantages over in-solution based digestion strategies. Firstly, throughput is vastly boosted to enable the efficient processing of up-to 96 samples in one step. Secondly, human introduced variability is curtailed to deliver more robust data. Thirdly, as protein is extracted from its lysis buffer before digestion, any detergent can be leveraged to boost protein denaturation pre-digestion, without compromising protease efficiency. As such, the comparable performance of in-solution vs SP3 digestion for rChymoSelect suggests that performance can be boosted by leveraging these denaturation strategies to optimize performance.

Varying urea concentration demonstrated no significant differences in protein identifications or sequence coverage distribution (Fig. 4e). However, manipulating guanidinium hydrochloride denaturant concentration and protease/protein ratio revealed more significant changes. An inverse relation between guanidinium hydrochloride concentration and protein identifications was observed, such that 0.1M boosted protein identifications by 5.6%, relative to 0.4M (Fig. 4f). Importantly, the functionality of rChymoSelect at 0.4M guanidinium hydrochloride demonstrates its applicability for proteins that require aggressive denaturation conditions, as was demonstrated with urea. As expected, increasing the protease/protein ratio also boosted identifications, with 1:20 driving increased protein identifications by 7.2%, vs 1:100 (Fig. 4g). We next probed whether increasing the rChymoSelect digestion time would drive a significant increase in protein identifications. For this, we compared a 2-hour incubation against a 16-hour incubation (with in-solution digestion, 2M urea and 1:40 protease:protein ratio). No significant change in protein identifications was observed when comparing the two conditions, with just a 2.4% increase from 2 hours to 16 hours (Fig. 4h). These data suggest rChymoSelect enables faster sample preparation, by avoiding the standardized overnight digestion, with minimal concomitant loss of identifications.

Following this, we investigated how this improved iteration of chymotrypsin would perform relative to trypsin, the gold standard of proteomics research. For this, we applied a deep proteomics pipeline, including offline peptide fractionation of lysed U2-OS cells before LC-MS/MS analysis. As expected, for proteins that were identified with a sequence coverage lower than 50%, trypsin outperformed rChymoSelect, in terms of number of proteins identified. Intriguingly, for more “digestible” proteins, who’s sequence coverage was greater than 60%; rChymoSelect demonstrated improved identifications, particularly in the case of >90% sequence coverage. Thus, while trypsin remains the enzyme of choice for global deep proteomics, rChymoSelect provides complementary coverage for proteins where near-complete sequence coverage is required.

### rChymoSelect facilitates improvements in Data-Independent Acquisition proteomics strategies

Recently, Data-Independent Acquisition (DIA) has become the premier strategy for high-throughput protein-identifications^**24**^. Briefly, whilst Data-Dependent Acquisition modes sequentially fragment the most abundant precursor ions, DIA performs fragmentation over a pre-defined window of *m/z*’s, scanning *m/z* bins across the precursor mass range, for each eluted chromatographic peak. This enables greater protein identifications, as well as curtailing issues of missing values across replicates. From our comparison of DIA vs DDA for rChymoSelect based proteomics, DIA facilitated a 45% increase in protein identifications, vs DDA, against standard K562 cell lysate (Fig. 5a). Because of the heterogeneity of sequence cleavage specificity of standard chymotrypsin; DIA data processing methodologies struggle to manage the unpredictable chimeric spectra that are generated from fragmentation; Therefore, we reasoned that the superior cleavage specificity of rChymoSelect would alleviate such issues, improving DIA workflows with chymotrypsin-based proteomics. Whilst a 3.6% increase in protein identifications was observed from a library-free DIA search with rChymoSelect vs standard chymotrypsin (Fig. 5b), the use of a DDA-generated spectral library within the DIA pipeline generated a 16.6% increase in peptide-spectrum matches, leading to a 4.6% increase in protein identifications (with 2M urea denaturation). Further, 4M urea denaturation increased PSMs and protein identifications by 22.4% and 6.2%, vs standard chymotrypsin, respectively (Fig. 5c, d).

**Fig. 5:**
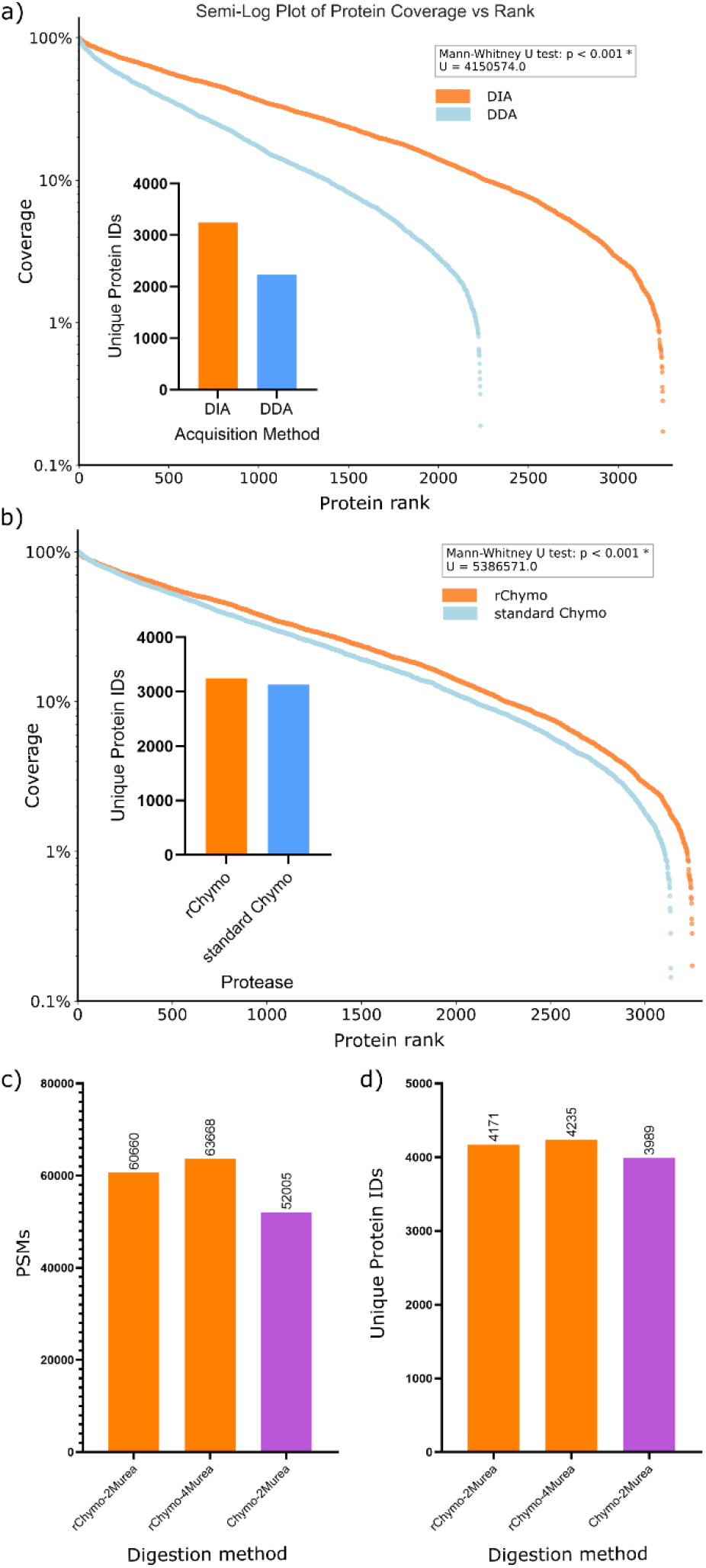
Incorporation of Data-Independent Acquisition (DIA) strategies into chymotrypsin proteomics. Protein sequence coverage distribution plot and unique protein identifications bar-chart for (a) for DIA vs DDA with rChymoSelect, and (b) rChymoSelect-DIA vs standard chymotrypsin-DIA, using standard K562 cell lysate. Samples were digested in 2M urea, using in-solution digestion, with 2-hour incubation at 25°C (library-free search). Spectral library-based DIA searches are represented in terms of (c) PSM identifications and (d) unique protein identifications.

Whilst these DIA experiments were performed against one experimental replicate per condition, the data are encouraging in the context of leveraging the advantages of rChymoSelect against the increased protein identifications, and robustness across replicates, that are associated with DIA strategies.

## Conclusions

Chymotrypsin offers a complementary analytical capability for proteomic applications through its preferential cleavage at hydrophobic amino acid residues, a specificity that is orthogonal to trypsin. Nevertheless, its broader substrate specificity and tendency to produce missed cleavages can limit digestion efficiency and expand the computational search space required for peptide identification during data processing. Benchmarking of a novel recombinant chymotrypsin analogue, rChymoSelect (Promega), against the conventional bovine chymotrypsin demonstrated marked improvements in cleavage specificity and proteome sequence coverage. Whereas standard chymotrypsin cleaved at the C-termini of nine distinct amino acid residues, the recombinant chymotrypsin variant exhibited vastly enhanced specificity, with up to 97% of peptide bond cleavages occurring after leucine, phenylalanine, or tyrosine residues. This increased cleavage selectivity led to fewer missed cleavages, reduced database search space, and resulted in a significant rise in peptide spectral matches, culminating in greater protein identifications and improved data reproducibility, relative to the standard enzyme. Further, the increased efficiency of proteolytic cleavage at hydrophobic residues facilitated greater recovery of mitochondrial and membrane-associated proteins, reflecting the elevated hydrophobic character of this cellular compartment. These findings place rChymoSelect as a valuable addition to the proteolytic enzyme toolbox, particularly for metabolism and membrane-focused proteomics experiments. Workflow optimization confirmed comparable performance between in-solution and SP3 digestion, supporting its flexibility in diverse sample-preparation formats. Further, denaturation with either 2M urea or 0.1M guanidinium hydrochloride provided sufficient protein unfolding for efficient proteolysis, although rChymoSelect’s effective functionality at up-to-6M urea and -0.4M guanidinium hydrochloride position it as highly attractive protease for the digestion of hydrophobic proteins that require harsh denaturation conditions. Increasing the incubation time from 2 hours to overnight (16 hours) did not yield a substantial improvement in proteome coverage, suggesting rChymoSelect to provide fast digestion efficiency, and enabling the entire sample preparation workflow to be completed in a matter of hours. Incorporation of Data-Independent Acquisition (DIA) strategies increased protein identifications by 45% relative to DDA, with reduced cleavage-site heterogeneity simplifying spectra and enhancing peptide assignments for rChymoSelect. Together, these findings establish rChymoSelect as highly valuable complementary protease for proteomics analysis.

## ASSOCIATED CONTENT

The mass spectrometry proteomics data have been deposited to the ProteomeXchange Consortium via the PRIDE^25^ partner repository with the dataset identifier PXD072165, Username, Password: ZY5JGOB1i6jX

## AUTHOR INFORMATION

### Author Contributions

The manuscript was written by K.R.A with contributions from all authors. All authors have given approval to the final version of the manuscript.

Conceptualisation of methodology: R. Z. C

Software: J. D Validation: K. R. A, J. D

Formal analysis: K. R. A, J. D

Investigation: K. R. A, J. D, G. H, R. Z. C

Writing-Original draft: K. R. A

Writing, editing and reviewing: K. R. A, K. T

Visualisation: K. R. A, J. D

Supervision: K. T, R. Z. C

Project administration: K. T, R. Z. C

Funding acquisition: K. T

## ABBREVIATIONS

rChymo: recombinant chymotrypsin
Chymo: chymotrypsin
SP3: Single-Pot, Solid-Phase-enhanced Sample Preparation
SDS: sodium dodecyl sulfate
SDC: sodium deoxycholate
GND: guanidinium hydrochloride
LC: liquid chromatography
MS: mass spectrometry
PSM: peptide spectral match
GO:CC: gene ontology:cell compartment
DDA: Data-Dependent Acquisition
DIA: Data-Independent Acquisition

